# Cognitive-and lifestyle-related microstructural variation in the ageing human hippocampus

**DOI:** 10.1101/2024.09.09.612084

**Authors:** Tyler Agyekum, Cindy L. García, Felix Fay, Olivier Parent, Aurélie Bussy, Gabriel A. Devenyi, M. Mallar Chakravarty

## Abstract

Ageing is a biological process associated with the natural degeneration of various regions of the brain. Alteration of neural tissue in the hippocampus with ageing typically results in cognitive decline that may serve as a risk factor for dementia and other neurodegenerative diseases. Modifiable lifestyle factors may help preserve hippocampal neural tissue (microstructure) and slow down neurodegeneration and thus promote healthy cognition in old age. In this study, we sought to identify potential modifiable lifestyle factors that may help preserve microstructure in the hippocampus. We used data from 494 subjects (36-100 years old) without clinical cognitive impairment from the Human Connectome Project-Aging study. We estimated hippocampal microstructure using myelin-sensitive (T1w/T2w ratio), inflammation-sensitive (MD) and fibre-sensitive (FA) MRI markers. Non-negative matrix factorization was used to integrate the signals of these images into a multivariate spatial signature of microstructure covariance across the hippocampus. The associations between hippocampal microstructural patterns and lifestyle factors & cognition were identified using partial least squares analysis. Our results reveal that the preservation of axon density and myelin in regions corresponding to subicular regions and CA1 to CA3 regions of the hippocampi of younger adults is associated with improved performance in executive function tasks, however, this is also associated with a decreased performance in memory tasks. We also show that microstructure is preserved across the hippocampus when there is normal hearing levels, physical fitness and normal insulin levels in younger adults of our study even in the presence of cardiovascular risk factors like high body mass index, blood pressure, triglycerides and blood glucose known to be associated with hippocampal neurodegeneration. This preservation is not observed in older adults when there are no normal levels of insulin, physical fitness and hearing. Taken together, our results suggest that certain lifestyle factors like normal hearing, physical fitness and normal insulin levels may help preserve hippocampal microstructure which may be useful in maintaining optimum performance on executive function tasks and potentially other modes of cognition.

## 1. Introduction

The hippocampus has long been known to play an important role in learning (Ranganath & Hsier, 2016), spatial navigation (Eichenbaum, 2017) and memory (Voss et al, 2017). It is also heavily implicated in the ageing process (Bussy et al., 2021) as it goes through atrophy throughout the course of the human lifespan (Betio et al., 2017; Bussy et al., 2021; R.S.C. Amaral et al., 2018) and, importantly, cognitive decline may be related to said atrophy (Betio et al., 2017; O’Shea et al., 2016; Murman, 2015; Fraser et al., 2015). This, of course, is a huge concern as the ageing population increases (Beard & Bloom, 2015).

Modifiable lifestyle factors are amongst the most promising avenues of promoting brain health during later stages in life (Livingston, 2017; Gottesman & Seshadri, 2022; Song et al., 2020). Positive factors like increasing physical activity, for example, have promoted neurogenesis in the hippocampus (Erickson et al. 2011; Sahay et al., 2011; Villemure et al., 2015). Cognitive stimulation in animal models has also promoted hippocampal neurogenesis (Betio et al., 2017) and caloric restriction without malnutrition potentially preserves cognitive function (Dahan et al., 2020). In contrast, negative factors like a higher body mass index and higher blood pressure have been associated with hippocampal atrophy (Raji et al., 2010; Cherbuin et al., 2015; Stranahan; 2015; Power et al., 2016). High fat and cholesterol diets resulted in an increase in microglia activation and phosphorylated-tau in the hippocampi of rats (Ledreux et al. 2016) and smoking and alcohol consumption also negatively correlate with hippocampal volume (Zahr et al. 2019; Nunes et al. 2019; Durazzo et al. 2014; Agartz et al. 1999).

The goal of this study is to identify how these modifiable lifestyle factors are associated with hippocampus structure as a putative window into the promotion of positive brain health. Unlike previous studies, we investigate associations with an under-examined dimension of brain anatomy in this context, namely the hippocampal microstructure. We estimated brain microstructure in vivo using magnetic resonance imaging (MRI) accessible measures (Patel et al., 2020; Patel et al., 2021; Robert et al.; 2022; Bussy et al., 2024) that are sensitive to various biophysical properties. In contrast to volumetric analyses, which fail to probe the variance of specific regional architecture in the brain (Patel et al., 2020). Measures obtained from diffusion MRI (dMRI) and from proxy measures that are correlated to myelin concentration are potentially sensitive to neuroanatomical variation that precedes classical volumetric changes (Callaghan et al., 2014; Lebel et al. 2008; 2018). Further still, these measures have been previously associated with cognitive variation (Charlton et al. 2013; den Heijer et al., 2012). As the hippocampus is a complex brain region with topographically-specific neuronal architecture (Amaral, 2018; Winterburn et al., 2013; Zeineh et al, 2017; Genon et al., 2021) there is evidence that microstructural properties in the hippocampus may be vulnerable to modifiable lifestyle factors (Treit et al., 2013; Lebel et al., 2018; Noble et al., 2013). For example, microstructure of white matter, have shown vulnerability to lifestyle factors like hypertension, obesity, diabetes, smoking, hearing loss, social isolation, depressive symptoms, and sleep disturbances negatively while physical activity, cognitive training, diet, and meditation have a positive effect on older individuals from the mid-life onwards (Wassenaar et al., 2019). Previously our group, used fractional anisotropy (FA), mean diffusivity (MD) and T1-weighted/T2-weighted ratio (T1w/T2w) to study the microstructure in both the hippocampus and the striatum (Patel et al, 2020; Robert et al., 2022). FA and MD are diffusion MRI derived metrics that have been known to describe axon geometry and axon density, respectively (Alexander et al., 2007; Jones et al., 2013; Tardif et al., 2016; Patel et al., 2020). T1w/T2w, on the other hand, is a ratio of T1-weighted images and T2-weighted images with signals that have been used as correlates for myelin density (Ganzetti et al., 2014; Glasser et al., 2014; Glasser & Van Essen, 2011; Grydeland et al., 2013; Tullo et al., 2019; Patel et al.; 2020; Robert et al., 2022). In isolation, these metrics are limited as they are not specific but sensitive to biological phenomena (Tardif et al., 2016; Jones et al., 2013; Patel et al., 2020), and their sensitivities to differing microstructural properties overlap with each other (Tardif et al., 2016; Jones et al., 2013; Patel et al., 2020).

To account for these challenges, our group has previously used orthogonal projective non-negative matrix factorization (OPNMF) (Patel et al, 2020; Robert et al., 2022) to integrate multiple neuroimaging measures in order to define optimal spatially ortoghonal dimensions of brain anatomy in a data-driven framework (Patel et al, 2020; Ochi et al., 2022; Robert et al., 2022; Patel et al., 2021).

In this study, we used OPNMF to investigate the relationships between hippocampal microstructure (defined as voxel-level measures of FA, MD and T1w/T2w) and modifiable lifestyle factors as well as cognition. First, we used OPNMF to define spatial microstructural patterns of neuroanatomical variance within the hippocampi of our cohort derived from the Human Connectome Project-Ageing (HCP-A) (Bookheimer et al., 2019). Next, we used these components to define inter-individual differences within these spatial patterns and related them to cognitive performance and modifiable lifestyle factors in separate analyses. We hypothesized that distinct hippocampal microstructural spatial patterns within our ageing cohort would be differentiated along the medial-temporal axis as has been defined classically in the hippocampus (Genon et al., 2021; Chauhan et al., 2021) and in previous work by our group (Patel et al., 2020), and their association with the variables related to healthy and maladaptive ageing, as in our previous work in the cortex (Patel et al., 2021; Bussy et al., 2024). We believe our results will allow us to understand potential patterns of lifestyle factors that may help preserve hippocampal microstructure and potential patterns of microstructure that are associated with cognitive performance.

## 2. Methods

### 2.1. Overview

A subset from the HCP-A dataset was selected for analysis (Data) alongside relevant demographic, cognitive, and lifestyle information. From HCP-A we used T1w and T2w images as inputs for the creation of a representative population template using multi-spectral nonlinear registration. Using the nonlinear transformations used to define a population-specific template. The T1w/T2w ratio, FA, and MD maps were warped into this same common space (Image Processing). Then, the whole hippocampus was segmented, and voxel-wise microstructural data was used as inputs for the OPNMF spatial decomposition.

The resulting subject-level OPNMF weights were used to describe microstructural inter-individual variability with neuroanatomical specificity (Orthogonal Projective Non-negative matrix factorization), which were then verified for generalizability (Stability Analysis). The relationship between these spatial dimensions of microstructural covariance and their relationship to cognition and modifiable lifestyle factors were then examined (Microstructure-cognition & Microstructure-modifiable lifestyle factors).

### 2.2. Data

We started with data from 719 healthy adults (age 36-100) drawn from the publicly available HCP-A dataset who had completed MRI acquisition across the measures of interest for this study (T1w, T2w, and diffusion) (Bookheimer et al., 2019). Sequence parameters have been previously published (Harms et al., 2018). We used T1w and T2w previously preprocessed through the HCP-A preprocessing pipeline (Glasser et al., 2013), which includes gradient distortion correction, alignment to MNI space, readout distortion correction, brain extraction, bias field correction and any repeated runs co-registered with 6 degrees of freedom (DOF) rigid body transformation (Glasser et al., 2013). Processed T2w images had also been registered to respective T1w images to create T1w/T2w image ratios (Glasser et al., 2013; Glasser & Van Essen, 2011).

494 (F/M: 286/208; Mean Age: 59.16 years; F Mean Age: 58.93; M Mean Age: 59.49; SD: 15.01; F SD: 15.32; M SD: 14.57 Handedness: 67.44; Race: 355 White, 66 Black, 38 Asian, 23 Multiracial, 10 unknown, 2 American Indian/Alaska Native) out of 719 subjects were analyzed for OPNMF due to 185 subjects missing cognitive and lifestyle factor data and 40 subjects failing quality control in the subsequent processing steps described below. Details of quality control can be found in the **Supplementary Materials** section 4.

### 2.3. Image Processing

Voxel-wise correspondence is required for the use of NMF to decompose the various input matrices. To enable this, we created a common space template (Patel et al., 2020; Robert et al., 2022) to which all microstructural maps (T1w/T2w, FA, and MD) were warped. First, a bounding box delimiting a distance of 10 mm from the hippocampus through all image dimensions was cropped for each subject’s T1w and T2w images. Hippocampi from all participants were segmented using the MAGeT Brain algorithm (Chakravarty et al., 2013; Pipitone et al., 2014). Following this, the ANTs multivariate construction tool (Avants et al., 2011) was used to create a population average using a multi-contrast optimization that used each subject’s cropped T1w and T2w. The nonlinear transformations that map each subject’s anatomy to the group average are then applied to each of the microstructural maps.

Hippocampus identification in the common space is achieved by applying the nonlinear transformations to MAGeT Brain hippocampus segmentations and which are then fused by majority vote as done previously in our group (Patel et al., 2020; Robert et al., 2022). Details for the model building and hippocampal segmentation are provided in **Supplementary Materials Section 1.1** and 1.2, respectively. Diffusion images were processed to obtain FA and MD maps using the FSL toolbox (https://fsl.fmrib.ox.ac.uk/fslcourse/2019_Beijing/lectures/FDT/fdt1.html) which included correction for susceptibility-induced artifacts with topup, eddy current correction, and fitting of the DTI model. Details for processing these data can be found in **Supplementary Materials Section 1.3**.

### 2.4. Orthogonal Projective Non-negative Matrix Factorization

To identify data-driven patterns of spatial covariance of the microstructure in the hippocampi of our population, we employed an orthogonal projective variant of NMF (OPNMF) (Sotiras et al., 2015; Yang & Oja, 2010). OPNMF provides a parts-based decomposition of input variables, which in our case was the voxel-wise microstructural scalar values of all subjects’ FA, MD and T1w/T2w images at the hippocampal level. OPNMF priortizes sparsity and orthogonal outputs, operationally provided components representing distinct non-overlapping spatial patterns. OPNMF decomposes an input matrix of dimensions m x n into a component matrix W (m x k) and a weight matrix H (k x n) where k is the number of components that needs to be specified by the user. Briefly, m takes on the dimension the number of voxels fused hippocampal label and n is 3 times the number of subjects (given the three microstructural measures). When OPNMF is run, the decomposition identifies k spatially distinct covariance patterns in W across subjects and metrics found in H. The final number of components used for downstream analyses is determined using a split half stability analysis and by examining the reconstruction error as proposed in our previous work (Patel et al., 2020, Robert et al., 2022; Kalantar-Hormozi et al., 2023; Patel et al., 2021). Details for the stability analysis and OPNMF implementation can be found in **Supplementary Materials Section 2**.

### 2.5. Microstructure-Lifestyle-Cognition Relationships

To understand the relationship between microstructure, lifestyle factors, and cognition we related individual microstructural covariance patterns (subject weights from OPNMF) with lifestyle factors and with cognitive performance using Partial Least Squares analysis (PLS) through Matlab version R2023a 9.14.0.2254940 with the Matlab PLS toolbox . The lifestyle factors included were: years of education (Livingston et al., 2020), friendship (Fortune et al., 2021), emotional support (Khondoker et al., 2017), stress (Bremner, 2006), sleep quality (Sextone et al., 2014), hearing (Chang et al., 2019), loneliness (Imai et al., 2022), peer rejection (Masten et al., 2009), metabolic equivalent of task (MET; Physical Activity) (Makizako et al., 2015), blood pressure (Korf et al., 2004), body mass index (BMI) (Cherbuin et al., 2015), cholesterol ratio (Dhikav and Anand, 2012), triglycerides (Nyberg et al., 2020), insulin (Biessels and Reagan, 2015), glucose (Kerti et al, 2013), and mineral electrolytes (Potassium, Sodium and Magnesium). For cognitive performance, we considered 9 variables in our studys; all of which have been linked to the various functions of the hippocampus. Details of these tests and respective cognitive domains are provided in **Supplementary Materials**, **Section 3.1.**

PLS is a multivariate technique that can be used to identify patterns of covariance between two different sets of data (Patel et al. 2020; Krishnan et al., 2011; McIntosh & Lobaugh, 2004; McIntosh & Mišić, 2013). The input data for the first PLS consisted of OPNMF subject weightings across the k components and the subject-wise results of the different cognitive test mentioned before. For the second PLS, the input data were OPNMF subject weightings at each of the k components and per-subject information regarding lifestyle factors. The PLS results consist of several latent variables (LV), that describe the covariance of hippocampal microstructure and cognition as well as the covariance of hippocampal microstructure and modifiable lifestyle factors. PLS also outputs the percentage of covariance explained by each LV.

To determine the statistical significance of each LV, permutation testing was used (10,000 permutations). And bootstrap resampling (10000 bootstraps) was used to estimate the standard errors and confidence intervals (Krishnan et al., 2011). More details can be found in the **Supplementary Materials Section 3.2.**

#### 2.5.1. Age and Sex analysis

Using RStudio 2023.09.1, we ran linear regressions to determine if there were age and/or sex effects and/or interactions on the microstructural scores and cognitive/lifestyle factor scores with the LVs obtained from the PLS analysis. Microstructural scores and cognitive/lifestyle factor scores were defined as singular values obtained from the decomposition of the microstructural and cognitive/lifestyle factor matrices used in the PLS analysis. The following linear model was used:

Microstructural scores or cognitive/lifestyle factor scores ∼ Age + Sex + Age:Sex

## 3. Results

### 3.1. Stability Analysis

To select the number of components, we analyzed both the stability and the accuracy, a description of this is provided in **Supplementary Materials** Section 2.1. **Figure 2A** and **Figure 2B s**how the stability plots for the stability coefficient (red) and the gradient of the reconstruction error (blue) from k = 2 to 10 for the left and right hippocampus respectively. In the left hippocampus, the stability coefficient plateaus at k ≥ 6. Peak stability coefficients were found for k = 4 and k = 2. In the right hippocampus, the stability coefficient has an inverse relationship with k until k = 4. From k = 4 until k = 6, there is a positive relationship between the stability coefficient k until it drops off and then levels out at k > 6. The results suggest that k = 4, k = 5, or k = 6 were all justifiable choices to use as a stable number of components for OPNMF. Meaning, after decomposition, there will be a k number of spatially distinct hippocampal patterns when either of those values are used.

Overall, there was a positive relationship between the gradient of the reconstruction error and k in both the left and right hippocampus. In the left hippocampus, there is a steeper change in the reduction of the reconstruction error moving from k = 4 to k =5 compared to the comparisons between the other k values and their adjacent. In a previous study by our group (Patel, 2020), k = 4 was chosen as the most suitable k value to use for further extended analysis of the hippocampus, and these analyses support this as a viable choice.

### 3.2. Neuroanatomical Description of 4-component solution

Figure 2 shows the results of an OPNMF 4-component solution for both the left and right hippocampus and a 3-dimensional volumetric rendering and representative coronal slices for the spatial maps of the OPNMF results. Figure 2E shows the subject weight matrix that describes the microstructural patterns of the subjects across all components at both a group and individual level. All subject weights were z-scored and shifted in this process to bring uniformity across the different weights in different components.

**Figure 1.**
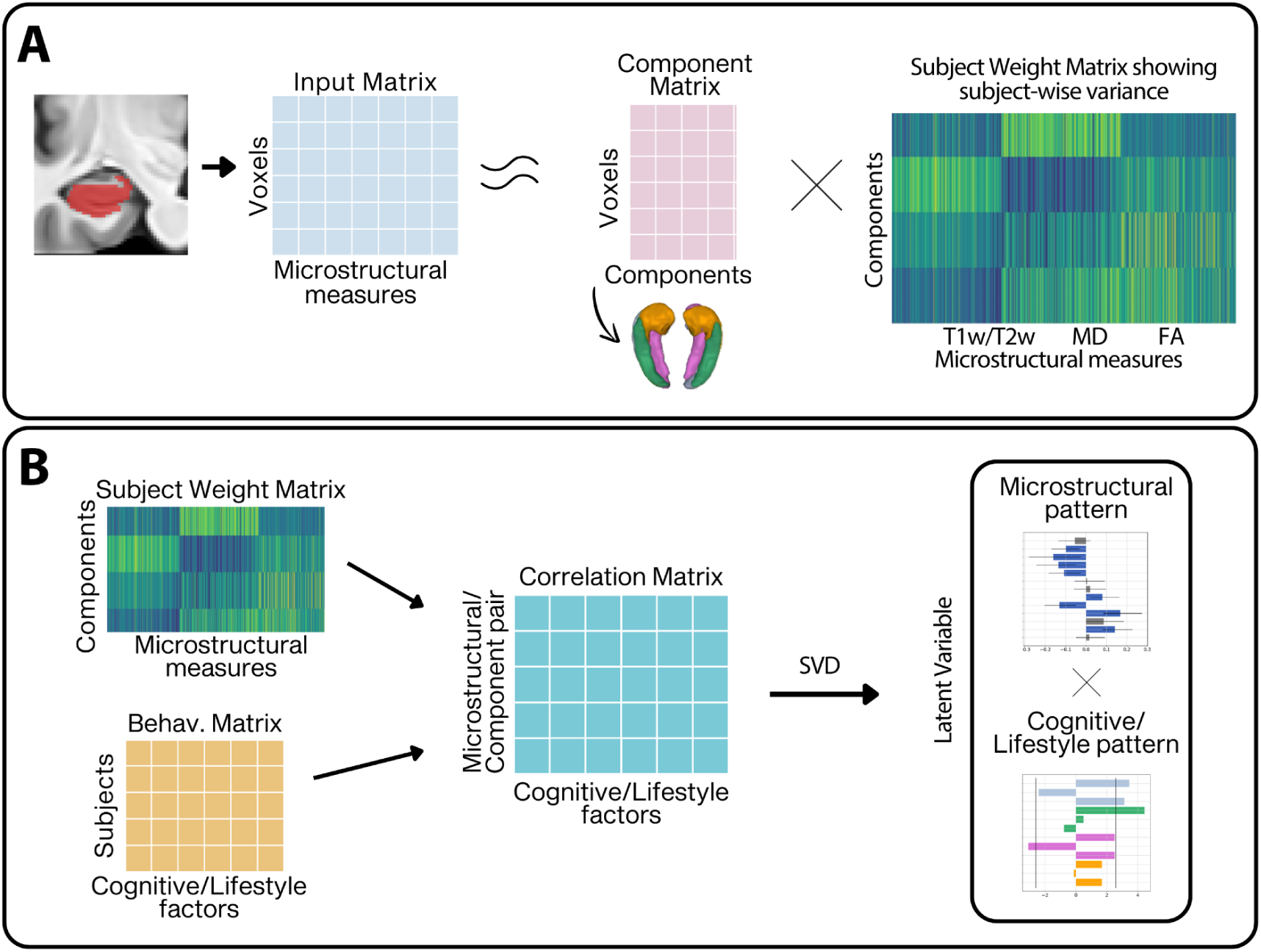
Workflow. A) We used automatically segmented hippocampal labels to extract our chosen microstructural metrics. We then concatenated hippocampal microstructural voxels into column vectors of our subjects to build an input matrix. Using orthogonal projective non-negative matrix factorization (OPNMF) we extracted spatially distinct patterns that represented the hippocampal microstructure covariance of our cohort. OPNMF decomposed the input matrix into a spatial component matrix and a subject weight matrix. B) Partial Least Squares Analysis (PLS) was then used to identify patterns of covariance between the subject weight matrix with cognitive variables and modifiable lifestyle factors each independently. This resulted in a set of Latent variables (LV) with both a microstructural pattern and a cognitive/lifestyle factor pattern.

**Figure 2.**
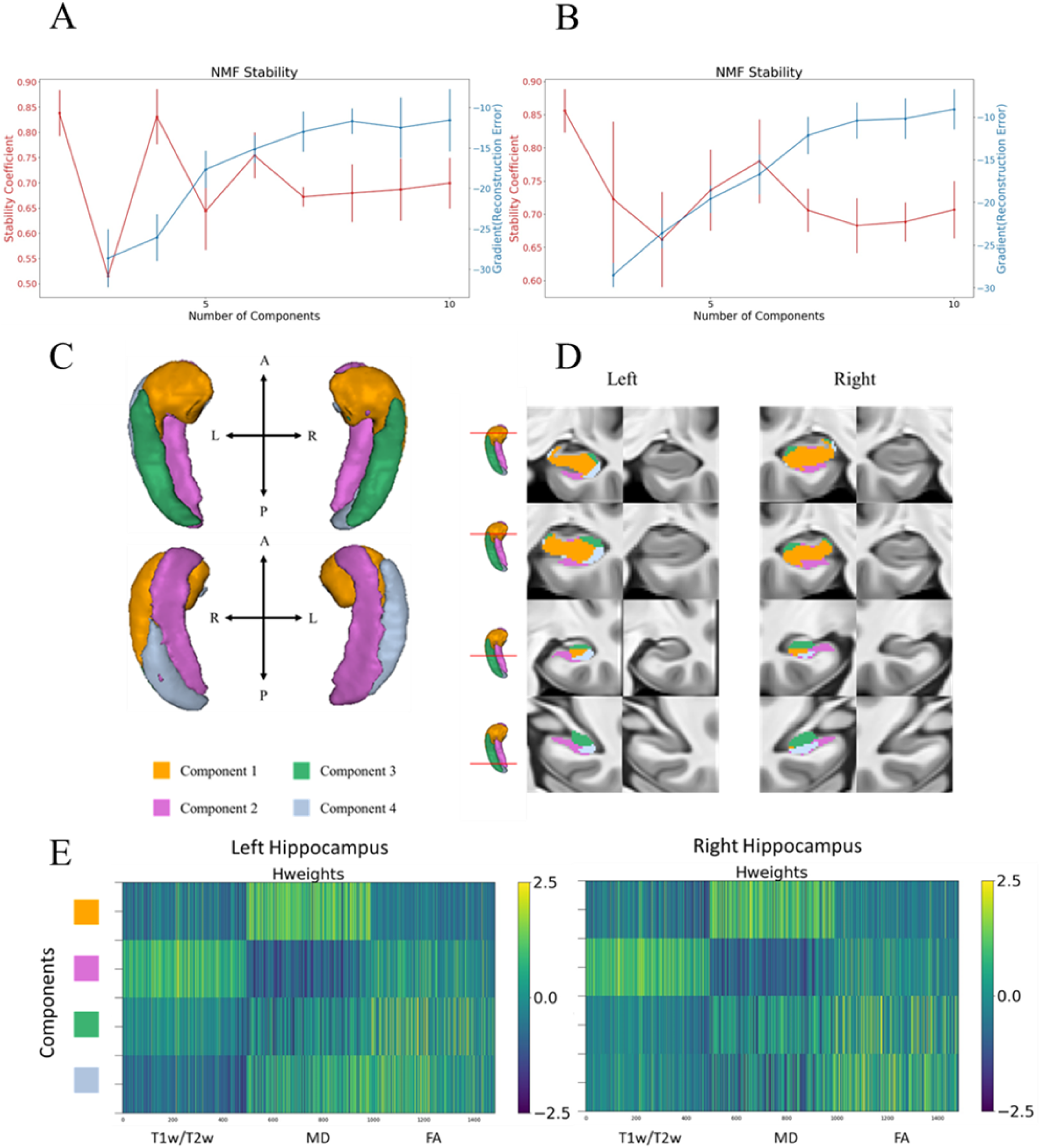
displays the stability plots in Figure 2A and Figure 2B, the OPNMF 4-component solution of the left and right hippocampus with a spatial map as both a 3-dimensional volumetric rendering in Figure 2C and coronal views in Figure 2D across the anterior-posterior axis, and the normalized subject weight matrix in Figure 2E.

**Figure 3.**
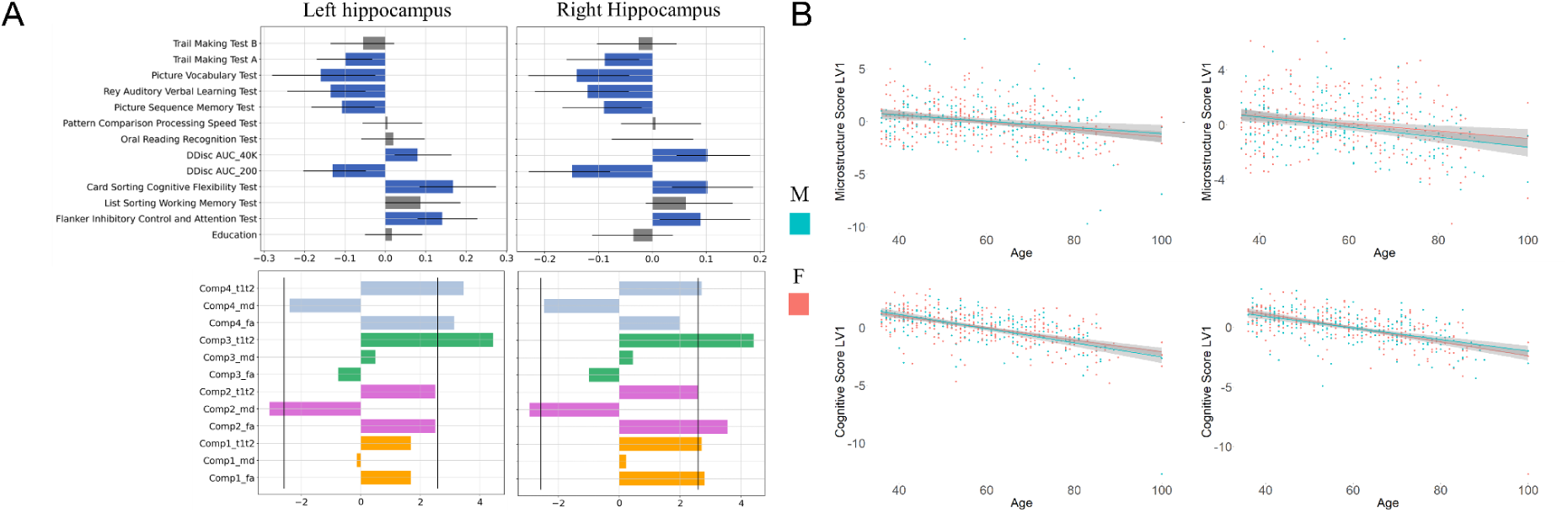
The Partial Least Squares output for cognition and microstructural patterns **(3A)** and the relationships between age and cognitive scores in both of the respective latent variables (LV) **(3B)**. Figure 3A shows cognitive (top row) and microstructural (bottom row) patterns of the left and right hippocampus. In the top row, the y-axis denotes the cognitive measure while the x-axis denotes the correlation of these measures with component-microstructural patterns of the same LV. In the bottom row, the y-axis denotes the component-microstructural metric while the x-axis denotes the bootstrap ratio. The black vertical lines denotate abootstrap ratio of ±2.58 (99% confidence interval), metrics surpassing this value are significantly correlated with the cognitive patterns in 3A. Figure 3B shows the relationship between age and the latent variable (LV) microstructure score in both brain hemispheres (top row) and the relationship between age and the LV cognitive score. The x-axes denotes the ages of the subjects in the study and the y-axes denote microstructure score (top row) and the cognitive score (bottom row).

**Figure 4.**
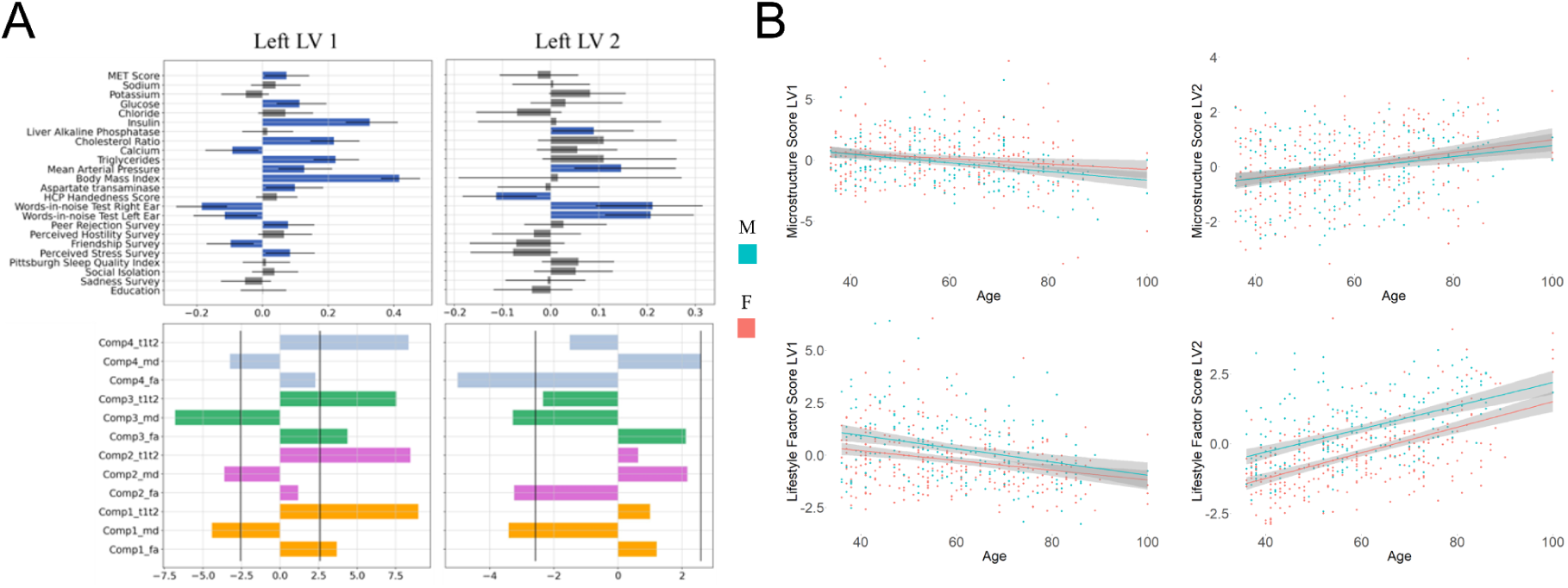
The left hippocampus Partial Least Squares output for modifiable lifestyle factors and microstructural patterns **(4A)** and the relationships between age and lifestyle factor scores in both of the respective latent variables (LV) **(4B)**. Figure 4A shows modifiable lifestyle factors (top row) and microstructural (bottom row) patterns. In the top row, the y-axis denotes the lifestyle factors while the x-axis denotes the correlation of these measures with component-microstructural patterns of the same LV. In the bottom row, the y-axis denotes the component-microstructural metric while the x-axis denotes the bootstrap ratio. The black vertical lines denote a bootstrap ratio of ±2.58 (99% confidence interval), metrics surpassing this value are significantly correlated with the cognitive patterns in 4A. Figure 4B shows the relationship between age and the latent variable (LV) microstructure score (top row) and the relationship between age and the LV lifestyle factor score (bottom row). The x-axes denotes the ages of the subjects and the y-axes denote microstructure score (top row) and the lifestyle factor score (bottom row).

**Figure 5.**
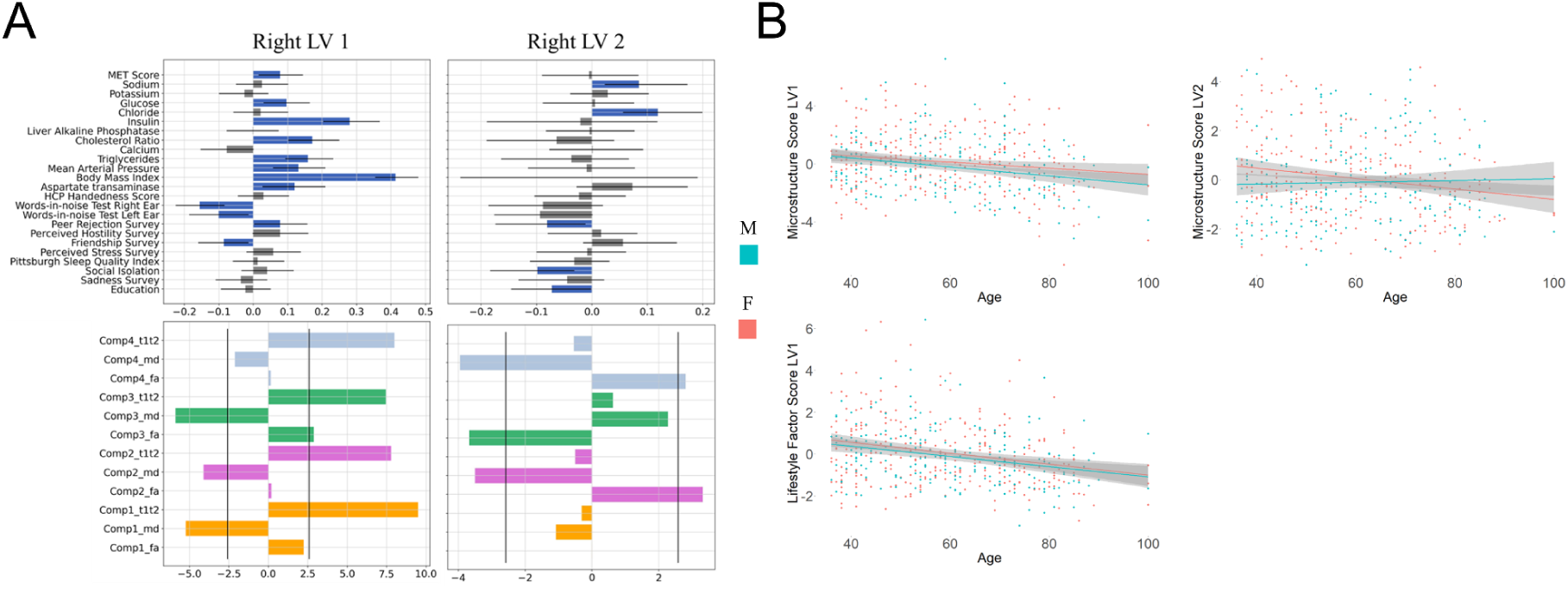
The right hippocampus Partial Least Squares output for modifiable lifestyle factors and microstructural patterns **(5A)** and the relationships between age and lifestyle factor scores in both of the respective latent variables (LV) **(5B)**. Figure 5A shows modifiable lifestyle factors (top row) and microstructural (bottom row) patterns. In the top row, the y-axis denotes the lifestyle factors while the x-axis denotes the correlation of these measures with component-microstructural patterns of the same LV. In the bottom row, the y-axis denotes the component-microstructural metric while the x-axis denotes the bootstrap ratio. The black vertical lines denote a bootstrap ratio of ±2.58 (99% confidence interval), metrics surpassing this value are significantly correlated with the cognitive patterns in 4A. Figure 5B shows the relationship between age and the latent variable (LV) microstructure score (top row) and the relationship between age and the LV lifestyle factor score (bottom row). The x-axes denotes the ages of the subjects and the y-axes denote microstructure score (top row) and the lifestyle factor score (bottom row).

Below, we describe each component using three main descriptors; its anatomical location along the anterior-posterior, lateral-medial, and superior-inferior axes within the hippocampus; its microstructural features; and how similar it is to previously defined hippocampal subfield locations.

**Component 1** is defined as having high MD, low FA, and low T1w/T2w (Figure 2E, row 1). It spans the whole anterior-posterior axis and is most prominent in the hippocampus head (Figure 2D, row 1). In the hippocampus head, it spans across the whole lateral-medial axis (Figure 2D, row 1). When looking across the superior-inferior axis, component 1 is mostly found in the superior portion of the hippocampus head, however, it is less present here when moving down the anterior-posterior axis (Figure 2D, rows 3 & 4). In comparison to hippocampal subfields, component 1 seems similar to CA4 and the dentate gyrus.

**Component 2** is characterized by low MD, high FA, and high T1w/T2w (Figures 2E, row 2). It also spans the whole anterior-posterior axis but in the head of the hippocampus it is more confined to the upper superior and lower inferior parts (Figure 2D, row 1 and 2). These two inferior and superior regions where component 2 is located are separated by component 1. Beyond the head of the hippocampus, component 2 is mostly confined to the inferior region of the hippocampus (Figure 2D, row 3 & 4). Across the lateral-medial axis, it is more confined to the medial portion of the hippocampus (Figure 2D, row 3 & 4). In comparison to the locations of hippocampal subfields, component 2 is found in a similar location to the subiculum, stratum and some parts of CA1.

**Component 3** is characterized by high FA, average MD, and averageT1w/T2w in the left hippocampus but a lower level of T1w/T2w in the right hippocampus (Figures 2E, row 3). Component 3 also spans the whole of the anterior-posterior axis; however, it is mostly confined to the superior region of the hippocampus (Figure 2D). In the head/more anterior region of the hippocampus, component 3 is also more confined to the medial and lateral portions of the hippocampus (Figure 2D, row 1 & 2). Below the head, component 3 is mostly confined to the lateral portion of the hippocampus (Figure 2D, row 3 & 4). In comparison to the location of hippocampal subfields, component 3 looks similar to CA3/CA2 regions.

**Component 4** is characterized by high FA in both the left and right hippocampi. T1w/T2w is averagein the right hippocampus but lower in the left hippocampus, and MD is high in the left hippocampus but average in the right hippocampus (Figures 2E, row 4).

Component 4 also spans across the anterior-posterior axis but is more prominent in the body-tail/posterior region of the hippocampus (Figure 2D, row 3 & 4). On the lateral-medial axis, component 4 is located more lateral to component 2, however, in the hippocampal head it is located in both lateral and medial areas (Figure 2D, row 1 & 2). It is also mostly confined to the inferior region along the superior-inferior axis. Finally, it is more prominent in the head region of the right hippocampus compared to the left hippocampus. Overall, the division of these components are similar to what was observed in the previous work by our group (Patel et al., 2020).

### 3.3. Microstructure-Cognition Relationships

Our PLS analyses, used to uncover interlinked dimensions of microstructure and cognition, demonstrated two significant LVs, one in the left hippocampus (ps < 0.001) and one in the right hippocampus (ps < 0.001).

#### 3.3.1. Left LV1

Left LV1 explained 64.85% of the covariance. The significant cognitive variables included decreased performance in the trail making A test (R = -0.09851 , 95% CI =[-0.1701, -0.03287]), decreased performance in the picture vocabulary test (R = -0.1602 , 95% CI = [-0.2807, -0.02523]), decreased performance in the Rey auditory verbal learning test (R = -0.1352, 95% CI = [-0.2435, -0.04943]), decreased performance in picture sequence memory (R = -0.1073, 95% CI = [-0.1838, -0.02700]), increased performance in delay discounting at $40,000 (R = 0.07925, 95% CI = [0.02271, 0.1630]), decreased performance in delay discounting at $200 (R = -0.1304 , 95% CI = [-0.2036, -0.04853]), increased performance in card sorting cognitive flexibility (R = 0.1676 , 95% CI = [0.08411, 0.2733]), and increased performance in Flanker inhibitory control and attention (R = 0.1410, 95% CI = [0.07913, 0.2273]). The associated microstructural features included increased T1w/T2w in component 3 and component 4, increased FA in component 4, and decreased MD in component 2.

#### 3.3.2. Right LV1

Right LV1 explained 71.98% of the covariance. The significant cognitive variables included decreased performance in the trail making A test (R = -0.0882, 95% CI = [-0.1598,-0.02484]), decreased performance in the picture vocabulary test (R = -0.1406, 95% CI = [-0.2307,-0.04278]), decreased performance in the Rey auditory verbal learning test (R = -0.1204 , 95% CI = [-0.2183, -0.04403]), decreased performance in picture sequence memory (R =-0.08973 , 95% CI = [-0.1674, -0.01992]), increased performance in delay discounting at $40,000 (R =0.1020 , 95% CI = [0.04445, 0.1808]), decreased performance in delay discounting at $200 (R =-0.1495 , 95% CI = [-0.2297, -0.07901]), increased performance in card sorting cognitive flexibility (R = 0.1018, 95% CI = [0.03584, 0.1867]), and increased performance in Flanker inhibitory control and attention (R = 0.08945, 95% CI = [0.01346, 0.1815]). Associated microstructural features included increased T1w/T2w in all components, increased FA in components 1-3, and decreased MD in component 2.

#### 3.3.3. Age and Sex analysis

A negative main effect of age on the microstructure score of LV1 in the left hippocampus was observed (p < 0.001, SE = 0.008502) and a similar effect was also observed in the right hippocampus (p < 0.01, SE = 0.008988). A negative main effect of age on the cognitive score of LV1 in the left hippocampus was observed (p < 0.001, SE = 0.005662) as well as in the right hippocampus (p < 0.001, SE = 0.005400).

#### 3.3.4. Summary

In both the left and right hippocampus, we observed a higher level of performance in cognitive flexibility, inhibition and attention, and delay discounting at 40k, alongside a lower level of performance in trail making test A, picture vocabulary, verbal learning and picture sequence memory. These patterns were associated with higher levels of T1w/T2w in component 3 & 4 and lower levels of MD in component 2. Bilateral effects occurred in relation to microstructure for FA in component 1 & 4 and T1w/T2w in component 4. Our age analysis showed that these patterns were more likely to be associated with younger individuals.

### 3.4. Microstructure-lifestyle Relationships

For the relationship between microstructural weights and modifiable lifestyle factors, our PLS results also revealed four significant latent variables, two in the left hippocampus (p < 0.001) and two in the right hippocampus; LV1 (p < 0.001), LV2 (p < 0.01).

#### 3.4.1. Left LV1

Left LV1 accounted for 78.46% of the covariance explained. The significant behavioural variables contributing to this LV included increased perceived sadness (R = 0.08338, 95% CI = [0.009286, 0.1587]), decreased perceived friendship (R = -0.09691, 95% CI = [-0.1701, -0.02702]), increased perceived peer rejection (R = 0.004093, 95% CI = [0.004093, 0.1577,]), decreased hearing loss in left ear (R = -0.1139, 95% CI = [-0.2103, -0.0157]), decreased hearing loss in right ear (R = -0.1846, 95% CI = [-0.2629, -0.1077]), increased aspartate transaminase (R = 0.09832, 95% CI = [0.009893, 0.1838]), increased body mass index (R = 0.4161, 95% CI = [0.3599, 0.4787]), increased mean arterial pressure (R = 0.1269, 95% CI = [0.0478, 0.2120]), increased triglycerides (R = 0.2221, 95% CI = [0.1555, 0.2929]), decreased calcium (R = -0.09213, 95% CI = [-0.1739, -0.01301]), increased cholesterol (R = 0.2175, 95% CI = [0.1453, 0.2948]), increased insulin (R = 0.3262, 95% CI = [0.2526, 0.4110]), increased glucose (R = 0.1126, 95% CI = [0.04332, 0.1941]), and increased MET (R = 0.07268, 95% CI = [0.006567, 0.1416]). The correlating microstructural features included increased T1w/T2w across all components, decreased MD across all components and increased FA in component 1 and 3.

#### 3.4.2. Left LV2

Left LV2 accounted for 10.39% of the covariance explained. The significant behavioural variables contributing to this latent variable included increased hearing loss in left ear (R = 0.2070, 95% CI = [0.1127, 0.2962]), increased hearing loss in right ear (R = 0.2112, 95% CI = [0.09311, 0.3144]), decreased HCP-A handedness score (R = -0.1133, 95% CI = [-0.1829, -0.02905,]), increased mean arterial pressure (R = 0.1460, 95% CI = [0.04870, 0.2586]), and increased alkaline phosphatase (R = 0.08925, 95% CI = [0.003231, 0.1719]). The correlating microstructural features included decreased FA in component 2 and 4, decreased MD in component 1 and 3, and increased MD in component 4.

#### 3.4.3. Right LV1

Right LV1 accounted for 77.43% of the covariance explained. The significant behavioural variables contributing to this latent variable included decreased perceived friendship (R = -0.08500, 95% CI = [-0.1603, -0.01401]), increased perceived peer rejection (R = 0.07792, 95% CI = [0.002327, 0.1569]), decreased hearing loss in left ear (R = -0.1010, 95% CI = [-0.1862, -0.014014]), decreased hearing loss in right ear (R = -0.1557, 95% CI = [-0.2254, -0.08325]), increased aspartate transaminase (R = 0.1201, 95% CI = [0.02699, 0.2090]), increased body mass index (R = 0.4128, 95% CI = [0.3542, 0.4786]), increased mean arterial pressure (R = 0.1312, 95% CI = [0.05742, 0.2092]), increased triglycerides (R = 0.1584, 95% CI = [0.09235, 0.2318]), increased cholesterol (R = 0.1712, 95% CI = [0.1012, 0.2495]), increased insulin (R = 0.2794, 95% CI = [0.2030, 0.3672]), increased glucose (R = 0.09678, 95% CI = [0.02944, 0.1637]), and increased MET (R = 0.07797, 95% CI = [0.015013, 0.14418]). The correlating microstructural features included increased T1w/T2w across all components, decreased MD across components 1, 2 and 3, and increased FA in component 3.

#### 3.4.4. Right LV2

Left LV2 accounted for 10.43% of the covariance explained. The significant behavioural variables contributing to this latent variable included decreased years of education (R = -0.07220, 95% CI = [-0.1458, -0.001588]), decreased perceived social isolation (R = -0.09820, 95% CI = [-0.1840, -0.03196]), decreased perceived peer rejection (R = -0.08073, 95% CI = [-0.1744, -0.01177,]), increased chloride (R = 0.1190, 95% CI = [0.05630, 0.1991]), and increased potassium (R = 0.08482, 95% CI = [0.02300, 0.1719]). The correlating microstructural features included increased FA in component 2 and 4, decreased FA in component 3 and decreased MD in component 2 and 4.

#### 3.4.5. Age and Sex analysis

A negative main effect of age on the brain score of LV1 in the left hippocampus was observed (p < 0.001, SE = 0.009353), however for the brain score in LV2 a positive main effect was observed (p < 0.001, SE = 0.005429). In the right hippocampus we also found a negative main effect of age on the brain score of LV1 (p < 0.001, SE = 0.009230). However, we did not find a main effect on age in LV2 but instead a positive main effect for females (p < 0.01, SE = 0.633930) and an interaction of age and sex (p < 0.05, SE = 0.01037) for brain scores. For behaviour scores with LV1 in the left hippocampus, we found a negative main effect for age (p < 0.001, SE = 0.006890) and a positive main effect for males (p < 0.05, SE = 0.5422). This negative main effect was also observed in the behaviour scores of the right hippocampus LV1 (p < 0.001, SE = 0.006796). For behaviour scores of LV2 in the left hippocampus, we found a positive main effect for age (p < 0.001, SE = 0.005084) and another positive main effect for males (p < 0.01, SE = 0.4000). However, there were no effects observed in the LV2 of the right hippocampus.

#### 3.4.6. Summary

In LV1 of both hemispheres, we observed that cardiovascular risk factors such as insulin, cholesterol ratio, triglycerides, BMI, positively correlate with T1w/T2w ratio across all components,but negatively correlate with MD across all components and positively correlate with FA in component 1 and 3. However, this pattern did not occur in the second latent variables when these positive factors were absent. Instead we see a pattern where there are reduced levels of FA in component 4 and 2 in the left and reduced levels of FA in component 3 in the right which both correspond to a lack of hippocampal microstructure preservation.

Our age and sex analysis also revealed that the microstructural patterns observed in LV1 of both hemispheres are more likely to be associated with younger individuals.

However, the microstructural patterns in left LV2 are more likely to be associated with older individuals. For right LV2, we observed an interaction showing that the microstructural pattern is more likely to be associated with older men and younger women. When it came to lifestyle factors, LV1 was more likely to be associated with younger individuals but left LV1 was more likely to be associated with males. Left LV2 was also more likely to be associated with males but with older individuals.

## 4. Discussion

In this study we used OPNMF to study microstructural covariance patterns of the hippocampus in an ageing cohort using FA, MD, and T1w/T2w. We identified microstructural patterns at both the group level and the subject level. Using PLS, we identified microstructural patterns that are associated with specific cognitive patterns and lifestyle patterns.

### 4.1. Group Level Microstructure

Previous work in our group (Patel et al., 2020; Robert et al., 2022), has shown the success of OPNMF at segregating complex regions of interest in the brain into meaningful patterns that are consistent with anatomical divisions. Below we discuss the hippocampal divisions observed in our work.

Our results showed component 1 having a higher magnitude of MD compared to FA and T1w/T2w while spatially shown to map out areas where the dentate gyrus (DG) and CA4 is located (Zeineh, 2017). We propose that the lower levels of FA and T1w/T2w that are associated with myelin (Alexander et al., 2007; Grydeland et al., 2013; Tardif et al., 2016; Ganzetti et al., 2014; Glasser et al., 2016), may be explained by the unmyelinated mossy fibres in these regions.

Component 2 showed a higher magnitude of FA and T1w/T2w compared to MD and the voxels were shown to correspond to subicular regions of the hippocampus (Zeineh et al., 2017). The subicular region of the hippocampus has been shown to have axon projections that arise from the entorhinal cortex known as the perforant pathway. Considering high FA has been associated with less fibre crossing, we posit that this pattern may be due to the direct fibre route of the perforant pathway in the subicular regions of the hippocampus.

In component 3, we saw a higher FA compared to MD and T1w/T2w and the voxels were shown to correspond to the CA2 to CA3 region of the hippocampus with reference to Zeineh and colleagues (2017). In these regions, schaffer collaterals have been shown to project from CA3 to CA2 (Zeineh et al, 2017) so it’s quite possible that this specific projection of axons may explain the high levels of FA. We also see lower FA and T1w/T2w in older participants and we posit that this may be due to the neurodegeneration of these collateral projections since lower FA and lower T1w/T2w have been associated with neurodegeneration (Alexander et al., 2007; Grydeland et al., 2013; Tardif et al., 2016; Ganzetti et al., 2014; Glasser et al., 2016).

Component 4 showed a higher FA and MD compared to T1w/T2w in the left hippocampus, however, in the right hippocampus, only FA was shown to be higher. This may have to do with the fact that the voxels of component 4 in the left hippocampus are prominent in the head region of the hippocampus but this is not true for component 4 in the right hippocampus. In reference to Zeineh and colleagues (2017) component 4 corresponds to the more inferior regions of CA1. As higher FA has been associated with less fibre crossing, the high FA in both hemispheres of the brain may be explained by the perforant pathway extending into the inferior parts of CA1 and the alvear pathway also extending through the inferior regions of CA1 (Zeineh et al., 2017). An ageing trend is also observed in both hemispheres with FA and T1w/Tw being lower in older adults and MD being higher. Based on our proposition, this may possibly be due to the pathways degenerating in older age. This view is supported by a post-mortem study where neurodegeneration was observed along the entorhinal-hippocampal pathway using FA as a microstructural metric (Uchida et al., 2023).

### 4.2. Individual Microstructural Variability

#### 4.2.1 Cognition and Hippocampal Microstructure

The improved performance on executive function tasks associated with the preservation of axon density in the CA1 to CA3 subicular regions may be explained by the connection between the hippocampus and the prefrontal cortex (PFC) through the entorhinal cortex as the PFC is associated with executive function tasks (Yuan & Raz, 2014; Buchsbaum et al., 2005; Rottschy et al., 2011) and the hippocampus has been implicated in this connection by retrieval suppression (Anderson et al., 2016). The decreased performance in other modes of cognition such as picture vocabulary, verbal learning, picture sequence memory, and trail making task B, however, were not consistent with previous studies where MD was inversely proportional to memory (den Heijer et al., 2012; van Norden et al., 2012).

The pattern for delay discounting associated with the preservation of hippocampal microstructure depended on the amount of money and this is consistent with the concept that the magnitude of the reward may play more of a role in delay discounting (Macedo et al. 2022).

The negative main effect observed in both hemispheres for both brain and behavioural scores shows us that the patterns observed here were more closely associated with the younger individuals in our study. These findings demonstrate the impact of age as a significant risk factor.

#### 4.2.2. Modifiable Lifestyle factors and Hippocampal Microstructure

The results for left and right LV1 show a preservation of microstructure across most of the hippocampus when there are significant levels of insulin, physical fitness and hearing in younger adults in our study despite the presence of cardiovascular risk factors like BMI, blood pressure, blood glucose and blood fat in younger adults. This concept has been shown in a number of animal studies (Maesako et al., 2012; Molteni et al., 2004). The preservation may be explained by insulin (Biessels & Reagan, 2015; Spinelli et al, 2019), physical fitness (Erickson et al, 2011; Sahay et al., 2011) and reduced hearing loss (Chang et al., 2019; Shim et al., 2023) being protective factors against atrophy the brain. This is further supported in left LV2 where an absence of these factors in the older adults of our study was associated with less microstructure preservation in the subicular and CA1 regions. Similarly, less microstructure preservation was observed in right LV2 around the CA2/CA3 regions when these specific lifestyle factors were absent.

#### 4.2.3 Age and Sex analysis

The preservation of microstructure in left and right LV1 being associated with the younger individuals of our cohort is consistent with the ageing trajectories in other studies (Mielke et al., 2012; Pereira et al., 2014; Bussy et al., 2021). Similarly, left LV2 more likely to be associated with older adults in our study is also consistent with ageing trajectories in other studies (Gunbey et al., 2014; Carlesimo et al., 2010). Older males and and younger females being more associated with the microstructural pattern in right LV2 showed a sex difference and this is consistent with previous studies that have demonstrated sex-specific brain ageing trajectories (Armstrong et al., 2019; Costantino & Paneni, 2020; Ruigrok et al., 2014).

The higher level of insulin associated with younger individuals in left LV1 is consistent with the concept of ageing having a deleterious effect on the secretion of insulin (Kurauti et al., 2021). We further posit that ageing may have played a role in how variables like insulin manifest and counteract negative variables. The finding of males showing a higher association with this pattern of modifiable lifestyle factors may be explained by sex-related differences in genes and hormones playing a role in different cardiovascular phenotypes (Regitz-Zagrosek & Gebhard, 2022). Older individuals being more likely to be associated with the negative lifestyle factors in left LV2 once again suggest that age is the primary risk factor and compounds the impact of other known factors such as blood pressure and hearing loss (Liu and Yan, 2007; Pinto, 2007). The left LV2 is also more likely to be associated with males in our study. As increased hearing loss is a part of this pattern, and there is a difference in hearing loss between sexes (Homans et al, 2016), this association lines up with previous studies. Other work has suggested this difference may be due to hormones (Shuster et al., 2019).

### 4.3. Limitations and Future Directions

One of the main limitations in our work is that the MRI accessible measures (T1w/T2w, MD, FA) used to tap into the microstructure of the hippocampus are sensitive but not specific to the biological phenomena that they measure. Various biological alterations at the brain tissue level ranging from myelination to neuroinflammation to changes in axon geometry may be detected by the MR signals but are not specific to them (Tardif et al., 2016; Jones et al., 2013; Patel et al., 2020). T1w/T2w, for example, while used as a correlate of myelin in our work and in previous works (Glasser et al., 2016; Glasser & Van Essen, 2011) has also been shown to be correlated with genetic markers and molecule size of oligodendrocytes and mitochondria (Ritchie et al., 2018). Whilst using three metrics compared to one metric may aid in characterizing the microstructure in our study, the above described limitations suggest that we can only use our interpretations of these measures to make hypotheses regarding the potential biological phenomena occurring at the tissue level. As we were limited by the MRI measures available in the HCP-A dataset, we suggest future studies consider using measures that may be more specific than that used in our study if available. Neurite orientation dispersion and density imaging (NODDI) for example, has shown promise to be a more specific marker of brain microstructure than FA (Zhang et al., 2012; Sacco et al., 2020).

Quantitative T1 mapping has also been used recently as a more specific marker for myelin (Marques et al., 2010; Marques & Gruetter, 2013; Sereno et al., 2013; Tardif et al., 2015). Nonetheless, we believe that the work presented here provides a robust framework for these future studies.

Another limitation is that the values of physical activity used to calculate MET scores were self-reported. Previous studies have explained that participants may respond to self-reported measures in a “normative” way for social desirability even when they are alone at home (Rosenman et al., 2011; Brenner & DeLamater, 2016). In this case, it is possible some measures used in our study may not reflect a participant’s actual level of physical activity. One potential option we suggest for future studies to measure physical fitness, is through the use of physical fitness trackers which has shown promise as a research tool (Bassett et al, 2019). Some of these devices have shown inter-device reliability for steps, distance, energy expenditure, and sleep (Evenson et al., 2015).

Finally, the data we used from the HCP-A dataset is cross-sectional and so we are limited by being unable to make any causal inferences from our results. As the HCP-A plans on eventually releasing a longitudinal subset of the cohort, it will be important to potentially replicate our work longitudinally. Specifically to see how some of the modifiable lifestyle factors have an effect on the hippocampal microstructure.

### 4.4. Conclusion

We used non-negative matrix factorization to analyze hippocampal microstructure by decomposing T1w/T2w, FA, and MD images in a multivariate manner. We further analyzed the relationship between hippocampal microstructure and cognitive variables and hippocampal microstructure and modifiable lifestyle factors using partial least squares analysis. This allowed us to identify hippocampal microstructural patterns that are associated with patterns of cognition and also identify modifiable lifestyle factor patterns that are associated with specific microstructural patterns. Specifically, we identified insulin, physical fitness, and reduced hearing loss as lifestyle factors that may help promote hippocampal health, particularly in middle aged adults.

