## Supplementary Materials for "Cognitive-and lifestyle-related microstructural variation in the ageing human hippocampus"

### Section 1: Image Processing

#### 1.1 Population Average Creation

A population average was created for the 719 subjects in the study to obtain voxel-wise spatial correspondence. First, a bounding box with limits of 10 mm from the hippocampus at all image dimensions and slices was cropped for each subject's T1w and T2w images. This helped improve registration at the area of interest and reduce computational expenses from registering the whole brain. Following this, the ANTs multivariate construction tool (Brian B. Avants et al., 2011) was used to create a population average using each subject's T1w and T2w images that had been cropped.  Both T1w and T2w images were used in the population average to improve registration by use of complementary contrasts.

Each subjects FA, MD and T1w/T2w images was then warped to the population average using antsApplyTransform with the population average serving as the reference image and b-spline interpolation serving as the method of resampling. Considering the population average was 0.8 mm isotropic in resolution in contrast to the 1.5 mm of the FA and MD images, this process upsampled FA and MD images to the population average. After warping, fslmaths was used to zero bound FA, MD and T1w/T2w images to remove any potential artifacts from b-spline interpolation. Filtering was applied to T1w/T2w images to remove potential outliers as described in previous studies (Glasser & Van Essen, 2011). In our case, however, geodesic distance was replaced by Euclidean distance considering our work involves volumetric space. This filtering involved the T1w/T2w value at a given voxel being compared to the 12 voxels around it in the x,y, and z direction; these 12 voxels were described as the given voxel's neighbourhood. Any T1w/T2w voxel value that was 2 standard deviations away from it's neighbourhood's mean was replaced with a gaussian weighted average of the neighbourhood (fwhm = 5 mm). Finally, hippocampal labels were warped to the population average using antsApplyTransform with GenericLabel resampling. Majority voting was used on all warped hippocampal labels that had passed MAGeT quality control to create one hippocampal label at template space.

#### 1.2 Automatic Hippocampus Segmentation

Subjects' hippocampus segmentations were obtained using the MAGeT brain algorithm tool (Chakravarty et al., 2013; Pipitone et al., 2014). This algorithm can segment a brain region by use of a multi-atlas procedure. Five manually segmented hippocampus atlases, including their white matter tracts, were used as an input for the algorithm (Amaral et al., 2016). Hippocampal subregions and white matter in these atlases include: cornus ammonis (CA) 1, CA2/CA3 (CA2CA3), CA4/dentate gyrus (CA4DG), stratum radiatum/stratum lacunosum/stratum moleculare (STRAT), subiculum (SUB), fimbria (FIMB), fornix (FNX), mamillary body (MB) and alveus (ALV) (Amaral et al., 2016). Twenty-one images from the subjects' T1w were selected as templates. The 5 atlases were non-linearly registered to the 21 subject templates and this created 21 cohort specific atlases. Non-linear registration of each template to each subject then created 105 atlas-template-subject candidates for each subject. Majority voting then occurred for each subject to get the final subject hippocampal segmentation. The minc-toolkit ANTs template construction tool (https://github.com/CoBrALab/optimized_antsMultivariateTemplateConstruction) was used for non-linear registration. Each hippocampus segmentation was then quality controlled using a protocol from our group (https://github.com/CoBrALab/documentation/wiki/MAGeT-Brain-Quality-Control-(QC)-Guide). Like previous work in our group (Patel et al., 2020; Sankar et al., 2017; Voineskos et al., 2015), hippocampus subfields were fused together to create one label per subject. Specific to this study, white matter tracts were removed and not included in subsequent analyses.

##### 1.3 Diffusion Preprocessing

We used unprocessed diffusion data from the HCP-A. Diffusion preprocessing was performed through the FSL diffusion toolbox to obtain FA and MD maps. This first involved using FSL topup (https://fsl.fmrib.ox.ac.uk/fsl/fslwiki/topup) (Andersson et al., 2003; Smith et al., 2004) to correct for susceptibility induced distortions and then FSL eddy (https://fsl.fmrib.ox.ac.uk/fsl/fslwiki/eddy) (Andersson and Sotiropoulos, 2016a) to correct for eddy currents and movements in the diffusion data. FSL eddyqc (https://fsl.fmrib.ox.ac.uk/fsl/fslwiki/eddyqc) (Bastiani et al. 2019) was also used to assess the quality of the data at a subject and group level. Finally, we used FSL dtifit to fit the diffusion tensor model to obtain scalar DTI maps of FA and MD.

#### Section 2: Orthogonal Projective Non-negative Matrix Factorization

For OPNMF analysis, the fused average hippocampal label for each of both the right and left hemispheres were used to extract FA, MD and T1w/T2w values using the TractREC package. These extracted voxels were stacked into a column of vector size (# of hippocampal voxels x 1 subject). This allowed us to obtain voxel-wise column vectors for each subject at each of the microstructural values (FA, MD, T1w/T2w). Consequently, this means each subject had 3 microstructural vectors per hemisphere of the brain leading to a total of 6 column vectors per subject. Input matrices were generated on a per-hemisphere basis and were of size voxels (NUMBER) by metrics and subjects (3 metrics * 494 subjects). After the input matrix was created, the data was normalized and then an OPNMF algorithm (Boutsidis and Gallopoulos, 2008; Halko et al., 2011; Sotiras et al., 2015; Yang and Oja, 2010) was applied to the input matrix using Octave 5.2.0. The OPNMF algorithm was initialized using non-negative double singular value decomposition (SVD) and used similar hypermeters as used in Patel et al. 2020 and Robert et al. 2022 (max iterations =100,000 and tolerance = 0.00001).

##### 2.1 Stability Analysis

To select the number of components, we analyzed both the stability and the accuracy. Analysis of accuracy was done by observing the gradient of the reconstruction error with an increasing iteration of the number of components (Patel et al., 2020; Durran, 1999; Fornberg, 1988; Quarteroni et al., 2007). We analyzed the stability by observing the similarities of spatial outputs over various splits of data. For example, our 719-sample size was randomly split into two (Wa = 359 & Wb = 360). This was done 10 times (k=1 to k=10) and then OPNMF was run on each split at each k independently. For each two splits, a cosine similarity between each row was calculated to compute a similarity matrix(c_Wa & c_Wb). Lastly, a coefficient ratio between each row was computed to find out if a given subset of voxels in one split corresponds with the same set of voxels in the other split. The mean correlation across all voxels informed us about the stability for a given granularity.

#### Section 3: PLS

##### 3.1 Cognitive tests

The cognitive tests considered were: dimensional change card sort test, that measures cognitive flexibility (Diamond et al., 2013), Flanker inhibitory control and attention test measuring inhibitory control and attention (Diamond, 2013), list sorting working memory test evaluating working memory, oral reading recognition test evaluating reading-decoding skills, pattern comparison processing speed test to measure processing speed, picture sequence memory test to measure episodic memory, Rey auditory verbal learning test to measure verbal episodic memory, picture vocabulary test evaluating subjects' vocabulary and the trail making test to measure processing speed. HCP standardized all scores to the National Institute for Health toolbox standard score for uniform analysis.

##### 3.2 PLS Statistical testing

The rows of the microstructural data were permuted 10000 times. Each permutation was used to compute a null distribution of singular values. The actual singular values were compared to this null distribution to derive p values for each LV. $P < 0.05$ used = 95% confidence that the singular value of non-permuted data was greater than that of the permuted data. This was done by random sampling where the rows in the behavioural matrix and the microstructure matrix was replaced randomly to create 1000 resampled sets of data, and this was then used to show a distribution of the singular vector of weight for each microstructural variable in each LV. The ratio of the singular vector weight over the standard error of the weight from the bootstrap distribution (bootstrap ratio, BSR) was used to determine the contribution and reliability of each microstructure variable. A BSR of 2.58 was used as this is analogous to a p value of 0.01 as used in other studies (Krishnan et al., 2011; McIntosh & Lobaugh, 2004; Nordin et al., 2018; Persson et al., 2014; Zeighami et al., 2017; Patel et al., 2020).

#### Section 4: Quality Control

At each stage of image processing (raw MRI motion artifacts, hippocampus segmentation, …), manual quality control (QC) was performed using the manual QC protocol that has been designed by our group (https://github.com/CoBrALab/documentation/wiki/Motion-Quality-Control-(QC)-Manual). Diffusion images were also further quality controlled through the automatic implementation of eddyqc from FSL (https://fsl.fmrib.ox.ac.uk/fsl/fslwiki/eddyqc).

**Figure 3.1** shows the population distributions of our starting subjects, those that passed QC after all processing steps, and those that failed QC.

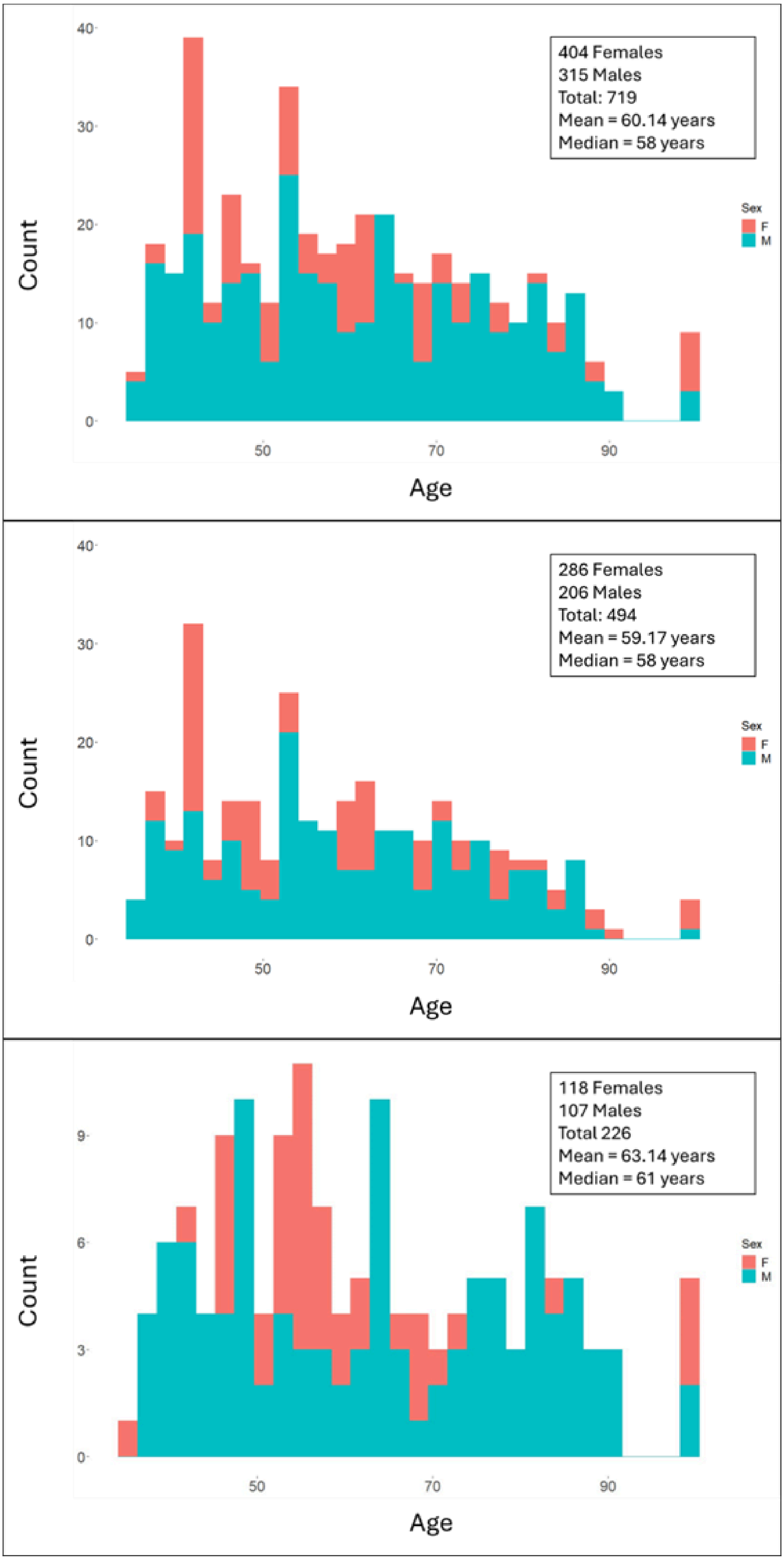
404 Females
315 Males
Total: 719
Mean = 60.14 years
Median = 58 years
Count
Age
Sex
F
M
286 Females
206 Males
Total: 494
Mean = 59.17 years
Median = 58 years
Count
Age
Sex
F
M
118 Females
107 Males
Total 226
Mean = 63.14 years
Median = 61 years
Count
Age
Sex
F
M

**Figure 3.1.** Population distributions of the starting subjects (top), subjects that passed QC (middle) and subjects that failed QC (bottom). X-axes show the ages of the subjects and y-axes show the amount of subjects within a given age. Subjects assigned female at birth are denoted by the pink-red colour and subjects assigned male at birth are denoted by the indigo colour.
